# Discrete genetic modules underlie divergent reproductive strategies in three-spined stickleback

**DOI:** 10.1101/2025.04.17.649467

**Authors:** Colby Behrens, Megan E. Tucker, Katie Julkowski, Alison M. Bell

## Abstract

A central challenge in biology is to understand how complex behaviors evolve. Given their importance to fitness, complex behavioral traits often evolve as an integrated package, but it is unclear whether suites of traits evolve through a few pleiotropic genetic changes, each affecting many behaviors, or by accumulating several changes that, when combined, give rise to an entire package of correlated traits. Typically, three-spined stickleback exhibit paternal care, a behavior that characterizes the entire *Gasterosteidae* family. However, an unusual “white” three-spined stickleback ecotype exhibits a suite of traits associated with the evolutionary loss of paternal care. In the white ecotype, males disperse embryos from their nests rather than care for them, build loose nests, exhibit high rates of courtship, and are relatively small in body size. These behavioral differences are apparent in stickleback reared in a common garden environment, suggesting the differences have a heritable basis. In an F2 intercross, we show that these traits are genetically uncorrelated and map to different genomic regions, suggesting that components of the white reproductive strategy segregate independently and evolved through the addition of multiple genetic changes. These results contribute to the growing body of evidence that the behavioral diversity observed in nature may often evolve by accumulating and combining alleles, each with modular effects, and show that this principle applies to a suite of behavioral traits that together form an integrated and adaptive strategy.

## INTRODUCTION

Behavioral traits are extraordinarily diverse in nature and often form suites of correlated traits that together comprise an integrated adaptive strategy. However, little is known about the underlying genetic basis of divergent behavioral strategies. Understanding how traits comprising a strategy are linked together, and how the strategies originally diverge, are outstanding questions that require mechanistic study. For example, it is unknown if novel behavioral strategies evolve via relatively few or many genetic changes (Mackay 2001; Mackay and Anholt 2024). Therefore, a comprehensive understanding of the genetic architecture of a trait, or suite of traits, could provide novel insights into the process of adaptation and speciation.

In many cases, genetic architecture of behaviors is examined using in-bred laboratory strains, which have been used to identify candidate genes and loci associated with a variety of behaviors (Flint 2003). However, there is a growing interest in examining natural variation of traits in wild populations (Greenwood et al. 2015; Jourjine & Hoekstra 2021). Indeed natural variation in nest-building, shoaling, and parental care behaviors has recently been examined in natural populations (Greenwood et al. 2015; Bendesky et al. 2017; Weber et al. 2013) to understand the genetic basis of complex behavioral traits such as those involved in social behavior. These studies often examine variation across species, but comparing behavioral strategies among recently diverged populations of the same species can offer insights into the initial proximate causes of divergence, before the lineages have accumulated differences that are confounded with the initial drivers that set the populations on different evolutionary trajectories.

The genetic basis of complex behavioral traits such as social behavior is challenging to study because they are typically influenced by many interacting genes of small effect and are sensitive to the environment. Modern sequencing technologies have enabled a variety of methods to identify genetic loci associated with behavior, including GWAS (genome-wide association studies), QTL (quantitative trait locus) mapping, transcriptomic profiling, and genome editing. In this study, we employ QTL mapping, which uses controlled experimental crosses (e.g. backcross, F2, recombinant inbred line) to associate phenotypic variation with genotypic variation. Such crosses can minimize background genetic variation and limit issues from population stratification (Broman 2001). Without considerable statistical power, QTL mapping usually cannot identify individual causal genes due to large confidence intervals, but QTL mapping with a moderate sample size (n∼100) may be able to identify general genomic regions which can enable inferences about the genetic architecture of a trait.

Here, we examined the genetic basis of divergent behavioral strategies between two ecotypes of three-spined stickleback fish (*Gasterosteus aculeatus*). Ordinarily, male stickleback provide paternal care to their developing offspring, but an unusual ecotype from Nova Scotia known as the “white” stickleback has evolutionarily lost paternal care (Blouw and Hagen 1990; Blouw 1996; Behrens et al. 2024). The “white” ecotype has recently diverged (< 12,000 years; Samuk 2016) from the sympatric “common” ecotype and the ecotypes differ in a variety of heritable morphological (coloration, size) and reproductive traits, including nest architecture, courtship behaviors, and paternal care (Blouw 1996; Behrens et al. 2024). Trait differences between the ecotypes are broadly consistent with sexual selection and parental investment theory (Kokko and Jennions 2008), where male whites are small, build weak nests whose primary function is to attract females, engage in high levels of female-directed courtship, and do not provide paternal care. In contrast, male commons are large, perform energetically demanding parental care, exhibit high levels of nest-directed courtship, and carefully build tidy nests for rearing their developing offspring. Together, the suite of traits contributes to distinct integrated behavioral and life history strategies for gaining fitness primarily via attracting more mates (whites) or producing viable offspring (commons). The common and white ecotypes often occur in sympatry and evidence of assortative mating is equivocal (Blouw and Hagen 1990; Corney and Weir 2023).

To determine whether the traits contributing to the white and common reproductive strategies are genetically separable, we generated a mapping population of F2 hybrids, phenotyped male behavior across major stages of the reproductive cycle (territoriality, nesting, courtship, parenting), and compared them to phenotypic variation in commons, whites and F1 hybrids (Behrens et al. 2024). We then performed quantitative trait locus (QTL) mapping to explore whether rapid divergence in reproductive strategies between the white and common ecotypes may have occurred via relatively few or many genetic changes. While our study lacks statistical power for fine mapping that would allow us to characterize the genetic architecture of these traits in detail, it represents an important first step toward understanding the genetic basis of divergence in reproductive strategies between the white and common ecotypes.

## METHODS

### Animal Husbandry

Adult F0 generation stickleback were collected from the wild in 2019. Common stickleback were collected from Cherry Burton Road, New Brunswick, CA (46° 01.516’N 64° 06.150’W), and white stickleback were collected from Canal Lake, Nova Scotia, CA (44°29’54.0”N 63°54’09.1”W). The fish were transported to the laboratory at the University of Illinois Urbana-Champaign and a single set of fish (grandparental generation) was artificially crossed to produce F1 hybrids, which were reared under standard conditions in the lab (Behrens et al. 2024) until they reached maturity in the summer of 2020. F1 hybrids (♀ Common x ♂ White; three families) were then mated in sib-ship crosses multiple times to generate a mapping population of F2 hybrids. To ensure the possibility for multiple clutches between the same individuals, F2s were generated through natural fertilizations. A gravid female was introduced to the male’s tank and allowed to interact for up to 15 minutes. Upon successful fertilization, the female and the eggs were immediately removed. The fertilized clutch was then placed in a mesh cup above an airstone (Day, Pritchard, and Schluter 1994), where it was artificially incubated; the embryos did not receive paternal care from their fathers. Once a mated F1 female became gravid again, she was reintroduced to the same male’s tank to generate additional clutches. Reproductive incompatibilities were not apparent in F2 hybrid offspring.

### Phenotyping

Upon maturation, signified by red throats and blue eyes (Wootton 1976), males were phenotyped following published methods (Behrens et al. 2024). We quantified traits at four stages of the reproductive cycle. Specifically, we measured territorial aggression, nest architecture, courtship, and parenting (specific behaviors are described in Table S1). Males were measured for standard length with digital calipers then moved from holding tanks to individual tanks (36L x 33W x 24H cm) containing nesting materials (filamentous algae, fake plant, sandbox, gravel). The males acclimated to the new tank for at least one day before experimental assays began. F0 and F1 trait variations were previously reported in Behrens et al. 2024 and are reproduced here. Experimental protocols were approved by the University of Illinois Urbana-Champaign IACUC (Protocol #18080 and #21031).

### Territorial Aggression

After building a nest, F2 male territorial aggression was measured with a “flask assay” (Huntingford 1976). Briefly, an intruder male stickleback was measured for standard length, size-matched to the focal male, and placed within a plugged Erlenmeyer flask. All intruders had signs of sexual maturity (red throats, blue eyes). The location of the focal male’s nest was recorded as being in one of three sections (left, middle, right). The observer waited 5 minutes to acclimate the focal male to their presence, then the flask was placed in the focal male’s tank in a section adjacent to his nest. If the nest was in the middle section, a coin toss was used to determine if the intruder was placed in the left or right section. Data collection began as soon as the focal male oriented towards the flask. We recorded the number of times the focal male bit against the flask and the time he spent oriented toward the flask with JWatcher (Blumstein, Evans, and Daniel 2006) for five minutes, after which the flask was removed from the tank. Additional courtship and nest-building behaviors (described in Behrens et al. 2024) were recorded but were too infrequent for QTL mapping. After 24 hours, the focal male was tested again with a random size-matched intruder, and the number of bites at the intruder was highly consistent across repeated trials (Kendall’s W = 0.455, n=133, p < 0.001).

### Nest Architecture

Nests were quantified according to a published scale that combined four major nest characteristics that are known to differ between ecotypes: nest height, sand content, nest opening, and nest location (Behrens et al. 2024). All traits were given an integer value from 1 to 5 and summed for a total nest score. Nest characteristics were quantified visually to avoid disturbing the male and were only quantified after a male exhibited nest-directed behaviors during a courtship trial.

### Courtship

Courtship behaviors were quantified in a free-swimming “no-choice” assays, as described in Behrens et al. 2024. Male and female behaviors were recorded for 15 minutes or until egg fertilization occurred. Male behaviors included zigzags, leads, bites, nest attendance, fanning, poking, and gluing (Table S1). If no fertilization occurred, the female was removed from the tank and the male was tested again after >24 hours. If fertilization did occur, the female was immediately removed and a parenting observation began.

Female whites were used in the courtship assays because clutches from female commons are stickier and more difficult to separate (Grant 1993), which could constrain males’ embryos dispersal behavior (Behrens et al. 2025).

### Parental Care

After fertilization, male parenting behaviors (Table S1) were recorded for 15 minutes. The frequency and timing of parenting behaviors were tracked with JWatcher (Blumstein, Evans, and Daniel 2006). The male was removed from the tank 30 minutes after fertilization and rapidly decapitated for tissue collection. During dissection, we quantified the number of cannibalized embryos present in the stomach and the number of embryos remaining in the nest as a proxy for future parental care.

### Tissue and DNA Extraction

Following behavioral trials, males were euthanized and tissues were dissected and saved for future analyses. Brains, eyes, kidneys, and testes were placed in RNALater. Livers were frozen over dry ice and stored without additional solutions. Fin clips were placed in ethanol. Muscle and skin samples were removed and placed in paraformaldehyde solution. Muscle was stored at 4°C and all other samples were stored at -80°C. DNA was extracted from fin clips with Qiagen Puregene Tissue kits, quantified via Qubit Broad-Range kits, and normalized across two 96-well plates.

### RADSeq and Genotyping

RAD library preparation was carried out by Floragenex. DNA samples were digested with the restriction enzyme PstI, fragments were size-selected to 300-500 bp, and sample-specific adaptors were ligated. The barcoded libraries were then pooled and sequenced on a NovaSeq6000 S4 lane with 2×150 bp reads, resulting in 5.3 billion reads. A total of 164 F2, 6 F1, and 2 F0 samples were submitted for sequencing. F2 samples were collected from 3 sib-ship families (n = 35, 26, 103). The F0 grandparents were sequenced twice to increase their sequence coverage.

Demultiplexing was performed using Stacks (Version 2.62; Catchen et al. 2013). Reads were aligned to the *Gasterosteus aculeatus* V5 reference genome (Jones et al. 2012; Peichel et al. 2020) using BWA (Li and Durbin 2009). We then followed the reference-aligned Stacks pipeline to identify SNPs with default filtering and r=0.8. Six individuals were dropped from analysis after alignment and Stacks due to low coverage (<3X). SNPs with fewer than 100 genotyped individuals were removed.

SNPs with the segregation pattern aaxbb were then imported to Lep-MAP3 (Rastas 2017) to build the linkage map. We used default setting with grandparent-based phasing unless otherwise noted. Briefly, SeparateChromosomes2 separated linkage groups with a minimum LOD of 22 and minimum linkage group size of 10. Unassigned markers were then added to linkage groups with the minimum LOD iteratively decreasing by 2 from 20 to 4. OrderMarkers2 then ordered the markers in each linkage group with 8 merge iterations and 8 polishing iterations. This ordering was performed 5 times with each linkage group and the highest likelihood order was saved. The final linkage map consisted of 21 linkage groups (20 autosomal, 1 sex) and 1853 markers across 1482 cM (Figure S1, Table S2).

### Statistical analysis

All statistical analyses were performed in R version 4.2.1 (R Developoment Core Team 2022). Traits from territorial aggression, nesting, and parenting assays were normal quantile rank transformed. Egg dispersal was treated as a binary variable.

Courtship data was collected across multiple trials for each male and therefore needed to be summarized at the individual level for QTL mapping. Since stickleback male courtship can change in intensity depending on how much interest is exhibited by a gravid female, we classified courtship trials into three groups: no interest, low interest, and high interest as in previous studies (Kozak and Boughman 2009; Behrens et al. 2024). To match previously reported behaviors from commons and whites (Behrens et al. 2024), we visualized F2 courtship behavior by plotting the first trial that a female exhibited low or high interest (Figure 1E-G). For QTL mapping, we tested for an effect of female interest level on F2 courtship behavior via a linear mixed model with female interest as a fixed effect and male identity as a random factor.

**Figure 1.**
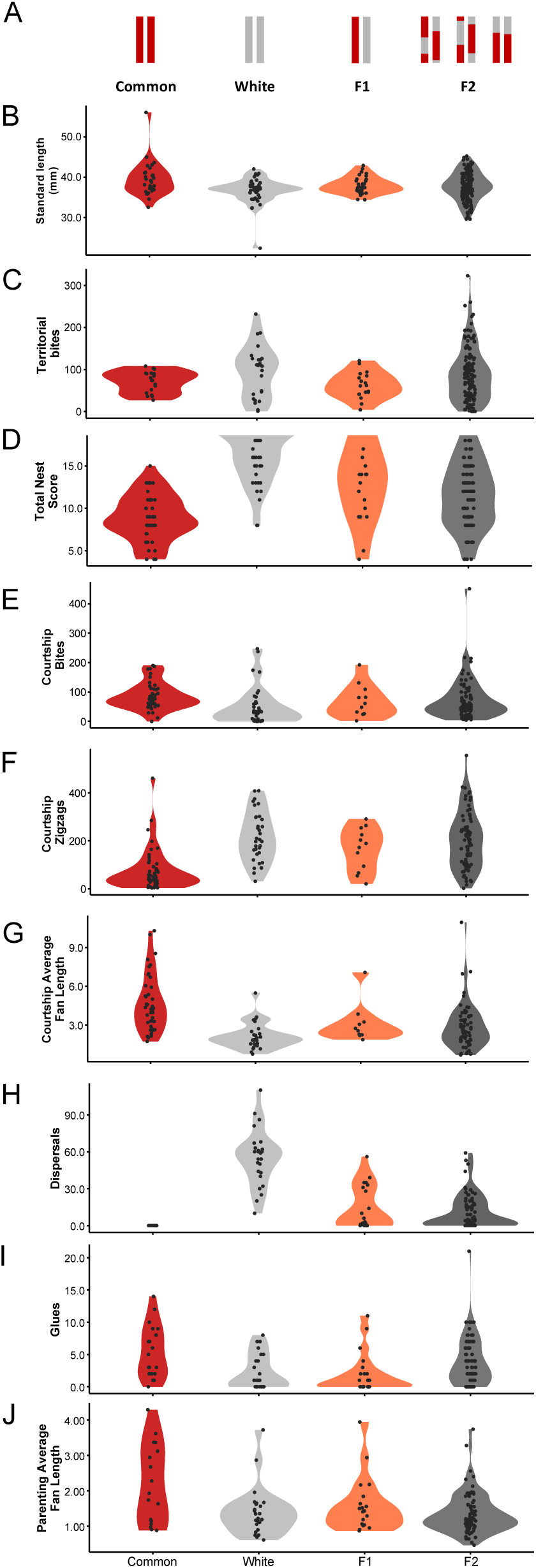
Distribution of reproductive behaviors in the stickleback ecotypes and their hybrids. (A) F1 hybrids were generated by crossing a female Common with a male White. Violin plots (B-J) display the distribution of behaviors across all four groups, with each dot representing a single individual. Data for commons, whites, and F1s are reprinted from Behrens et al. 2024.

Traits that were not associated with female interest were averaged across trials, while traits that were associated with female interest were Z-score normalized then averaged across trials for each male (Table S3).

Trait correlation matrices were compared using Box’s M test (*boxM*) from the package heplots (Fox, Friendly, and Monette 2009). Visualizations were created using the package corrplot (Wei et al. 2017).

QTL mapping was performed in the R package R/qtl (Broman et al. 2003). Markers placed at the same location on the map were moved slightly with the jittermap function. We performed mapping with Haley-Knott regression and determined significance thresholds through 1000 permutations of the data with the “scanone” function. Sample sizes ranged from n=76 to 133, so we characterized peaks with p < 0.1 as suggestive QTL (Greenwood et al. 2013). We calculated variance explained by the QTL with fitqtl, which estimates the percent variance explained by the peak marker of a QTL. Additionally, we scanned for pairwise interactions across the genome using the function “scantwo”. Significance thresholds were determined via 1000 permutations.

## RESULTS

### Genetic architecture of reproductive behaviors

We phenotyped 133, 121, 118, and 76 F2 males across the territorial, nesting, courtship, and parenting stages of the reproductive cycle, respectively. The F2 hybrids exhibited wide phenotypic distributions that encompassed variation observed in the common and white F0s (Figure 1), and the unimodal distribution of the traits suggested more than one locus is associated with each trait. To investigate whether the traits share a common genetic basis, we compared phenotypic trait correlations in the F0 and F2 populations. Traits were generally highly correlated in the F0 generation, but these correlations were reduced or lost in the F2 mapping population, especially across stages (Box’s M = 182.11, df = 91, p < 0.001; Figure 2), suggesting that the traits are genetically separable.

**Figure 2.**
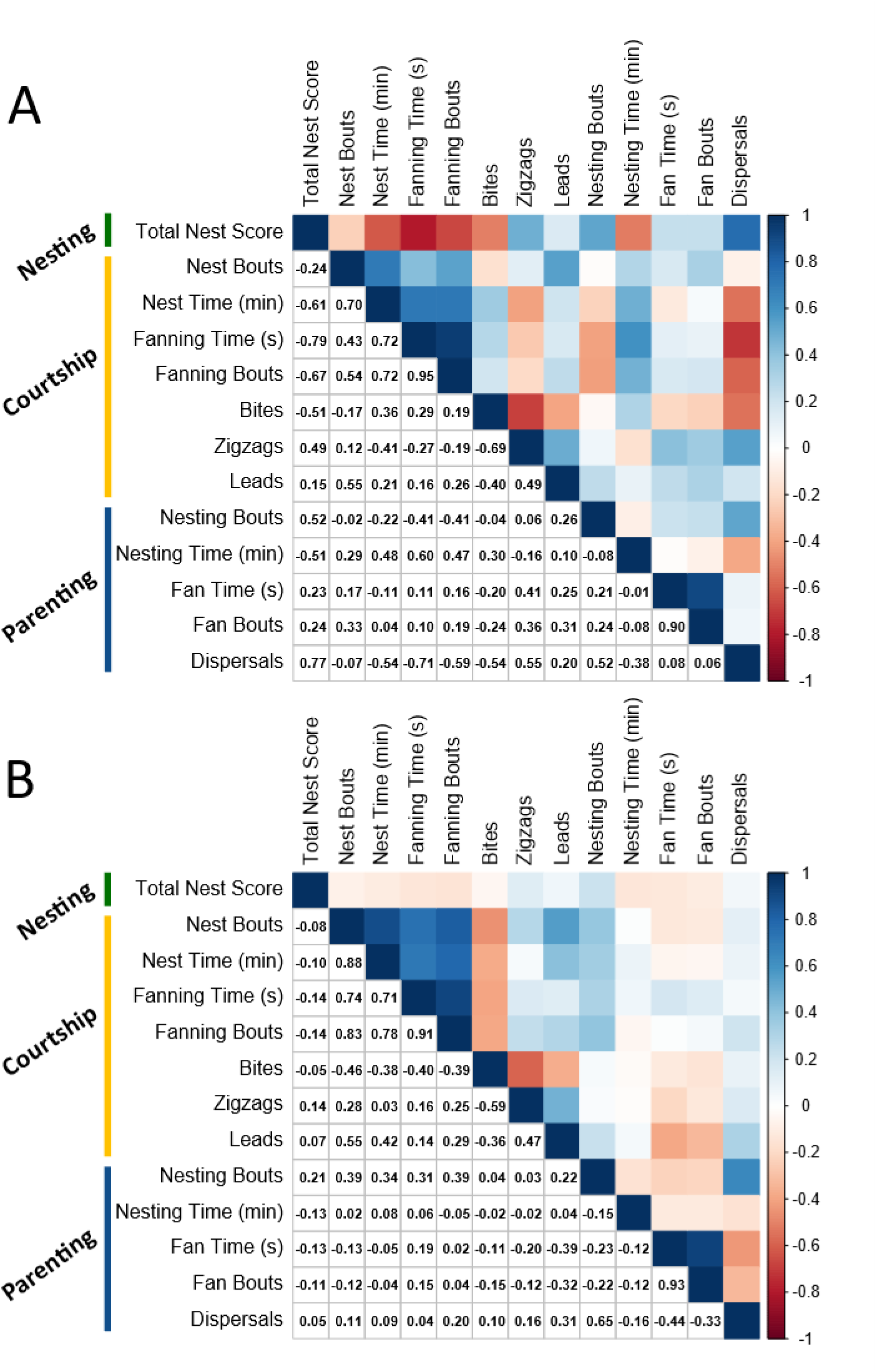
Trait correlations among F0 (A) and F2 (B) individuals. Three trait categories (nesting, courtship, parenting) are shown for each group, with the value of the Spearman correlation coefficient below the diagonal and a heat map of the value of the correlation coefficients above the diagonal (darker values indicates stronger correlations). The strength of correlations is generally low in the F2 generation, especially across behavioral categories, consistent with the hypothesis that the traits are genetically independent.

We then genotyped the F2s via RADseq, assembled a linkage map with the resulting 1,853 markers (Figure S1), and performed QTL mapping on traits of interest. The markers were evenly distributed at high densities across the genome (Table S2).

### Body size

Body size and morphology frequently vary across three-spined stickleback populations. In the populations studied here, commons are often larger than whites in the wild (Blouw and Hagen 1990) and when reared under similar conditions (Behrens et al. 2024; Behrens et al. in press). We detected a QTL significantly associated with standard length on chromosome 18 (LOD=4.94); individuals with homozygous common or heterozygous genotypes were larger than homozygous whites, matching patterns exhibited in natural populations (Figure S2). Additionally, two-dimensional scans detected an additional suggestive QTL on chromosome 6 (LOD = 9.35, p = 0.089, Figure S3A) that interacts with the locus on chromosome 18. Previous studies have detected QTL associated with body size on chromosomes 1, 6, 17, 18, 19, and 21 (Greenwood et al. 2011; Peichel and Marques 2017). and these QTL overlap with previously identified QTL associated with standard length (Chr 18; Greenwood et al. 2011) and body size (Chr 6; Liu et al. 2014) in stickleback.

### Territorial aggression

Territoriality is vital to stickleback reproduction, and males frequently bite and chase intruders and neighboring males to defend their nesting territory (Wootton 1976; Huntingford 1976). Whites, commons, and F1 hybrids did not significantly differ in aggression in response to a simulated territorial intrusion, though whites trended towards higher aggression (Common μ = 70.9 (SD 25.5), White μ = 94.8 (SD 61.1) bites; Figure 1C). We detected a QTL on chromosome 1 (LOD=4.67) that was significantly associated with territorial bites against an intruding male (Figure 3; Figure S4). Individuals that were homozygous for the white allele exhibited higher bite rates overall (Figure S4B), matching the pattern in the F0 populations. A recent study identified a QTL associated with territorial aggression on chromosome 16 in a cross between divergent stickleback populations in Japan (Yamazaki et al. 2025), but we did not see any evidence for that QTL in this study.

**Figure 3.**
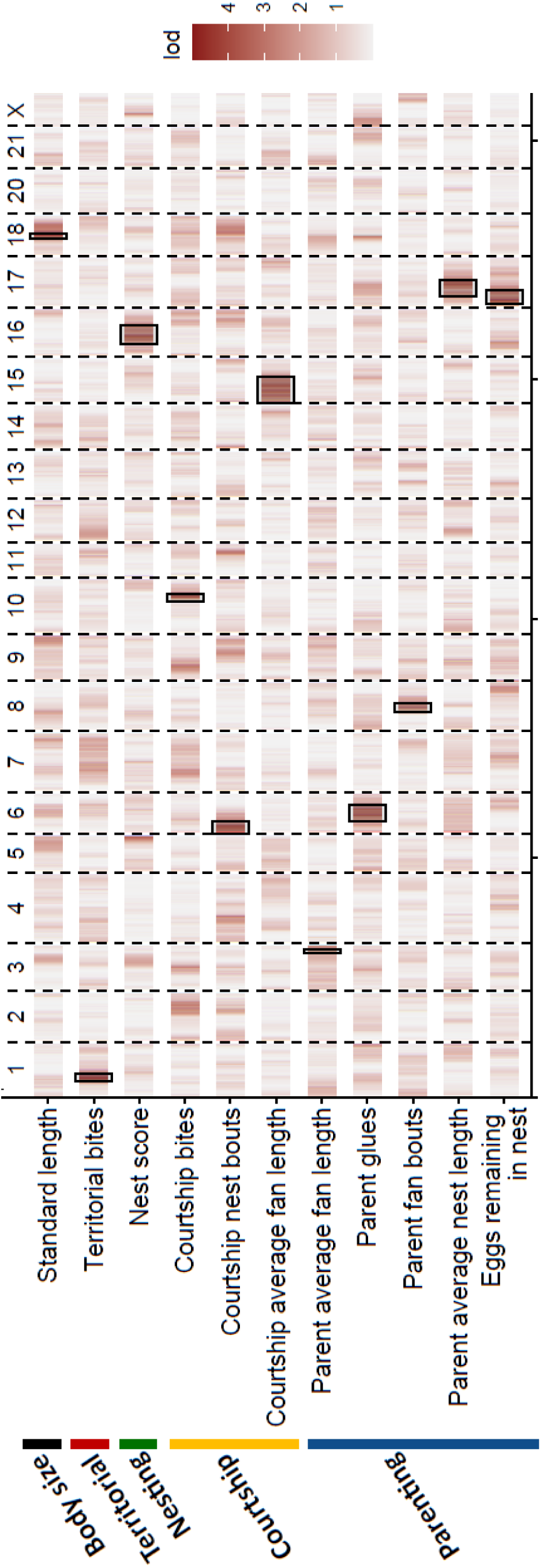
Reproductive behavior QTL (p<0.1) are distributed across the genome. The LOD score for each trait is plotted across all chromosomes, with darker sections indicating higher LOD scores. Significant peaks and their confidence intervals are indicated with boxes. Chromosome 19 is the sex chromosome and is renamed here as “X”.

### Nest architecture

Typically, nests are important for attracting potential mates (Barber, Nairn, and Huntingford 2001) and for rearing offspring in stickleback. Nests may also act as a prezygotic reproductive isolating mechanism if there are large differences in architecture or microhabitat (Blouw 1996; Behrens et al. 2024; Dean et al. 2021). Males of the common ecotype typically build flat, compact nests made of sand and algae with an opening on the top, while male whites build simpler, taller nests without sand with an opening on the side (Behrens et al. 2024). This variation can be quantified as a score on a 20 point scale incorporating sand, flatness, nest opening, and location; whites build significantly higher scoring nests (e.g. no sand, not flat, horizontal opening, within algae mass) than commons (Behrens et al. 2024). F2 hybrids exhibited extensive variation in nest architecture (Figure 1D), and we detected a QTL on chromosome 16 (LOD=3.8) that was significantly associated with nest score in F2 hybrids (Figure 3; Figure S5A). At this locus, the white allele was associated with a greater total nest score, matching the pattern in the F0 populations (Figure S5B). Other high, though non-significant, peaks were detected on chromosome 5 (LOD=3.24) and the X chromosome (LOD=2.76), suggesting a polygenic basis and possible sex linkage.

### Courtship behavior

Common and white stickleback differ in a variety of courtship behaviors. White stickleback perform more female-directed displays (zigzags, leads), while common stickleback perform more nest-directed displays (nest attendance, fanning, glues; (Behrens et al. 2024; Haley et al. 2019). After controlling for female receptivity, we detected QTL associated with three different courtship behaviors: average fanning length, nesting bouts and bites (Figure 3).

During courtship, commons often exhibited higher rates of nest-directed behaviors than whites (Figure 1G; Behrens et al. 2024). We detected separate QTL associated with two nest-directed behaviors: average fanning length (Chromosome 15) and nesting bouts (Chromosome 6; Figure 3; Figure S6). During courtship, commons engage in longer fanning bouts than whites, potentially as a signal to the female of future parental care (Behrens et al. 2024), and in the F2s, individuals with the homozygous common alleles at the QTL peak exhibited longer fanning bouts (Figure S6B).

Though bites are often interpreted as aggressive behaviors (Sevenster 1961), recent evidence suggests that they can also be important for signaling during courtship (Behrens et al. 2024). During courtship, male commons bite females significantly more than male whites do (Figure 1E), and female commons positively respond to bites more frequently than female whites (Behrens et al. 2024). In F2s, we detected a significant QTL on chromosome 10 (LOD=3.79) that is associated with biting during courtship. Though biting is mechanically similar during territorial defense and courtship, territorial and courtship bites mapped to separate chromosomes (chromosome 1 versus 10). These data suggest that distinct sets of genes are involved in regulating the same motor pattern that serves different functions and that timing or context of execution of the motor pattern, not the motor pattern itself, has evolved in whites.

Standard genome scans did not detect any QTL associated with several of the courtship behaviors that strongly differentiate common and white stickleback (e.g. zigzags and leads) and which are expected to have a genetic basis (Blouw 1996; Behrens et al. 2024). However, two-dimensional genome scans detected suggestive interacting loci associated with courtship leads. Individuals with homozygous common alleles at marker sites on chromosomes 6 and 16 exhibited fewer leads (LOD 9.76, p = 0.055, Figure S3B), again suggesting that the observed behavioral variation has a complex genetic architecture that cannot be fully explained by individual genes or loci.

### Parental behavior

Parental care is typically necessary for offspring survival in stickleback (van Iersel 1953; Blouw 1996). In most populations, male stickleback fan, glue, and attend the nest after fertilizing eggs. White stickleback have evolutionarily lost paternal care (Blouw 1996); instead of providing care to the developing offspring, male whites remove embryos from their nests and disperse them into the surrounding area (Figure 1H). In addition, male whites exhibit significantly reduced parental behaviors like gluing (Figure 1I) and fanning (Figure 1J). In F2 hybrids, parenting behaviors are highly variable (Figure 1H-J) and we detected five QTL that were associated with different parental behaviors: glues, fan bouts, average fanning length, average nesting length, and the number of eggs remaining in the nest (Table 1).

**Table 1.** Locations of significant (p<0.1) QTL. The 11 QTL are distributed across 9 chromosomes.

| Trait category | Trait | Chr | cM | Marker | LOD | pval | PVE | Interval Size (Mb) |
| --- | --- | --- | --- | --- | --- | --- | --- | --- |
| Body size | Standard length | 18 | 36 | c18.loc36 | 4.94 | 0.008 | 13.24 | 4.89 |
| Territorial | Bites | 1 | 34.00 | 7311 | 4.67 | 0.008 | 13.71 | 1.27 |
| Nesting | Total nest score | 16 | 30.97 | 732074 | 3.8 | 0.082 | 12.14 | 11.09 |
| Courtship | Nest bouts | 6 | 10.64 | c6.loc10 | 3.93 | 0.047 | 12.63 | 2.47 |
| Courtship | Bites | 10 | 60.00 | c10.loc60 | 3.79 | 0.063 | 12.22 | 1.29 |
| Courtship | Average fan bout length | 15 | 27.42 | 685060 | 3.87 | 0.063 | 12.46 | 14.96 |
| Parenting | Average nest bout length | 17 | 40.15 | 780958 | 3.86 | 0.069 | 19.3 | 12.66 |
| Parenting | Glues | 6 | 29.62 | 344016 | 3.82 | 0.083 | 19.1 | 12.02 |
| Parenting | Fan bouts | 8 | 35.88 | 442425 | 3.72 | 0.084 | 18.64 | 3.47 |
| Parenting | Average fan bout length | 3 | 67.38 | 140428 | 3.62 | 0.092 | 18.59 | 1.07 |
| Parenting | Eggs in nest | 17 | 10.00 | c17.loc10 | 4.17 | 0.042 | 22.06 | 1.94 |

When building and maintaining nests, male stickleback apply a glycoprotein glue (spiggin) to their nests (Kawahara and Nishida 2007; Seear et al. 2015). The rate of gluing varies across time and contexts, suggesting that the glue may have a function beyond nest maintenance (Kozak, Head, and Boughman 2011; Behrens et al. 2024), and commons apply more glue immediately after fertilization, indicating a potential role in early parental care. We detected a QTL associated with gluing on chromosome 6 (LOD=3.82) in F2 hybrids (Figure S6). Consistent with the effects of natural selection, the allelic impact matched expected patterns (Figure S6B), as the common allele at this locus led to greater rates of gluing in F2 hybrids. This QTL – along with others with allelic patterns that matched the expected patterns from the F0 generation (e.g., body size (SL), territorial bites, total nest score, courtship fanning, parenting glues) – may be good sites for future genetic dissection.

Males fan their nests both to attract mates and to provide parental care, and male commons perform more fanning behavior in both contexts than male whites. Interestingly, the average fanning length performed in the courtship context mapped to a different chromosome than the average fanning length performed in the parental context (chromosome 15 versus 3, specifically), again suggesting that distinct sets of genes are associated with the same motor pattern in different contexts.

Two parental behaviors (average nesting bout length and eggs remaining in the nest) mapped to overlapping QTL intervals on chromosome 17. After fertilization, commons engage in longer nesting bouts than whites (Behrens et al. 2024). Similarly, commons do not remove eggs from their nest, while whites disperse their embryos (Blouw 1996; Behrens et al. 2024). Eggs remaining in the nest can therefore be used as a proxy for potential future parental care. Despite strong differentiation between commons and whites, we did not detect significant QTL associated with dispersal behavior from one- or two-dimensional genome scans. However, two-dimensional scans detected a potential, though non-significant, interaction between markers on chromosomes 17 and 20 (Figure S3C). Individuals with homozygous common alleles at both sites performed few egg dispersals, matching the natural patterns in F0 populations (Figure 1H). Additionally, the presence of an overlapping QTL for all three traits suggests a common genetic basis for the divergence in parenting behaviors.

### QTL associated with divergent behaviors are widespread across the genome

After mapping, we identified one QTL associated with a morphological difference (standard length) and 10 QTL associated with variation in behaviors at the territorial, nesting, courtship, and parenting stages (Figure 3, Table 1, Figure S8). The QTL associated with behavioral traits were distributed across nine chromosomes, suggesting that these behaviors often have distinct genetic bases. Each QTL explained a significant proportion of the phenotypic variation (12.14 – 22.06%) though these effect sizes are likely overestimates due to the limited sample size (Würschum and Kraft 2014) and the confidence intervals are wide, so likely include many genes.

## DISCUSSION

Here, we examined the underlying genetic architecture of reproductive behaviors between a pair of stickleback populations that differ in whether they provide paternal care. F2 hybrids exhibited extensive phenotypic variation across a myriad of behavioral traits related to breeding, correlations among traits were uncoupled in the F2 mapping population, and QTL associated with divergent traits were widely distributed across the genome. Together, these results suggest that multiple behavioral traits comprising the divergent breeding strategies are genetically independent. According to genomic comparisons, overall genome divergence between whites and commons is low (genome-wide F_ST_ ∼0.02) and the populations are differentiated by small F_ST_ outlier regions across the genome (Samuk 2016). Therefore, several lines of evidence suggest that the white ecotype evolved via multiple genetic changes, each contributing to distinct components of their reproductive strategy.

These consistent results beg the question: how did the highly divergent white reproductive strategy evolve so quickly (<∼12,000 years; Samuk 2016)? Other examples of evolutionary divergence in deer mice (Hager et al. 2022), finches (Funk et al. 2021), *Drosophila* (Durmaz et al. 2018), and Atlantic cod (Berg et al. 2017), have implicated chromosomal rearrangements or inversions with the rapid evolution of a suite of traits, but considering that key divergent traits studied here mapped to different genomic regions, this does not seem to apply in the white three-spined stickleback. In other populations of stickleback, standing genetic variation in marine populations has been important for facilitating rapid adaptation to freshwater environments (Colosimo et al. 2005; Jones et al. 2012). It is possible that strong selection on standing genetic variation contributed to rapid divergence between the white and common reproductive strategies in these Atlantic Canadian populations as well. Stickleback populations in the Outer Hebrides in Scotland also differ in a variety of nesting and courtship traits (Dean et al. 2021) and potentially differ in parental care (personal correspondence), and another loss of parental care has been reported in stickleback in the Baltic (Borg 1985). Further study of convergent evolution in these additional populations could examine the role of standing genetic variation in facilitating the independent evolutionary loss of care in replicate stickleback populations. Additionally, the black-spotted stickleback (*Gasterosteus wheatlandii*), the closest living relative to three-spined stickleback, exhibits egg manipulation that is qualitatively similar to the egg dispersal seen in the white ecotype (McInerney 1969). This prompts another (nonexclusive) hypothesis, which is that the genetic variation contributing to the white strategy is very old, potentially predating divergence between *wheatlandii* and *aculeatus*, estimated at 14 million years (Varadharajan et al. 2019).

A major goal of evolutionary biology is to identify the loci, and genes, associated with phenotypic diversity. Here, we detected QTL associated with a variety of traits (morphological and behavioral) throughout the stickleback reproductive cycle. In several cases, the effects of QTL were biologically meaningful. For example, F2 hybrids with homozygous white alleles on chromosome 16 built nests that closely resembled nests built by male whites. These results bolster a wealth of studies on the genetic basis of ecologically relevant traits in three-spined stickleback (Peichel and Marques 2017) by supporting known QTL (e.g. body size QTL on chromosomes 6 and 18) and by identifying new genetic associations. The size of QTL intervals were variable (1.07 to 14.96 Mb) and could contain hundreds of genes (supplemental data), so integrating these candidate genes with other lines of evidence, including regulatory information from RNA expression studies, could provide additional insights into the genetic basis of these traits.

Though we detected several QTL associated with rapidly evolving behaviors of interest, we did not detect individual QTL associated with some of the genetically-based traits strongly differentiating commons and whites (e.g. courtship zigzags, courtship leads, post-spawning egg dispersal; Behrens et al. 2024). It is likely that lack of statistical power to identify loci of small effect contributed to the failure to identify QTL associated with these traits. Behavioral traits are often associated with QTL of small effect (Flint 2003; MacKay et al. 2009), and the power to detect a significant QTL is closely associated with both the cross design and size of the mapping population (Broman 2001). Our F2 mapping population was relatively small, particularly in later reproductive stages, so courtship and parenting traits were therefore more difficult to map.

The traits examined here also likely have a highly polygenic genetic architecture, as suggested by the largely unimodal trait distribution in F2 hybrids. Indeed, two-dimensional genome scans suggested interacting QTL for two behaviors, courtship leads and egg dispersals, that were not identified with one-dimensional scans. In both cases, an individual required homozygous common alleles at both sites to exhibit behaviors similar to a common male. Another limitation that may have contributed to the failure to identify loci is if some of the traits are Y-linked, and thus not mappable in this study because we included just one cross direction (♀ Common x ♂ White). Some white and common behaviors are thought to be sex-linked (Behrens et al. 2024), and a closer examination of the sex chromosomes in this system could provide insights into their divergence

Finally, it is possible that there are significant QTL associated with these behavioral traits, but they were not detected due to limitations of the phenotyping methods and/or their environmentally and socially sensitive nature. This could be especially true for traits involved in social interactions, such as courtship behavior, that are very plastic. Males often alter their courtship behavior, for example, in response to female behavior (Guevara-Fiore, Stapley, and Watt 2010; Cole and Endler 2016), and in stickleback the frequency and proportion of courtship behaviors are highly dependent on female receptivity (Behrens et al. 2024). Variation in female interest was accounted for statistically in this study, but variable levels of female interest and receptivity may have obscured genetic signals and made association mapping for traits involved in courtship more difficult.

Gene association mapping studies have pointed to the importance of discrete genetic modules in natural behavioral variation; for example, the tendency to school and body position when schooling are genetically dissociable in sticklebacks (Greenwood et al. 2013) and entrance tunnel length and presence of an escape tunnel are genetically independent in *Peromyscus* mice (Weber, Peterson, and Hoekstra 2013). Here, we provide evidence that the same principle applies to a male reproductive strategy which includes nest architecture, courtship behavior, and parental care; genes contributing to these components of male white and common reproductive strategies are genetically separable. Importantly, the success of the white strategy depends not only on males’ behavior, but also on coevolutionary changes in their mates and offspring. In particular, it benefits females to alter their parental investment in response to differences in male parental care; indeed, female whites produce relatively larger clutches of smaller eggs that are less adhesive than common females (Grant 1993; Behrens et al. in press). Moreover, white offspring have presumably adapted to no longer require extensive paternal care (sensu Jarrett et al. 2018a; Jarrett et al. 2018b). White offspring are more bold as juveniles (Neumann and Bell 2023), but little else is known about coevolutionary changes in offspring. Whether discrete genetic modules contribute to an even larger set of traits exhibited by other members of the family is unknown; answering that question has the potential to provide deep insights into the processes that contribute to the extensive diversity of family life among organisms.

### Statement of Authorship

C.B. and A.M.B conceptualized the project. C.B., M.T., and K.J. collected data. A.M.B secured funding and supervised the project. C.B. performed analyses and data visualization. C.B. and A.M.B wrote the original draft. C.B., M.T., K.J., and A.M.B reviewed and edited the manuscript.

## Acknowledgments

We thank Floragenex for sequencing services and Anne Dalziel and members of the Bell lab for providing valuable comments and feedback. Experimental protocols were approved by the University of Illinois Urbana-Champaign IACUC (Protocol #18080 and #21031).

## Data and Code Availability

Code and data for reproducing the QTL analyses and visualizations will be made available at Dryad.

## Supplemental Figures

**Figure S1.**
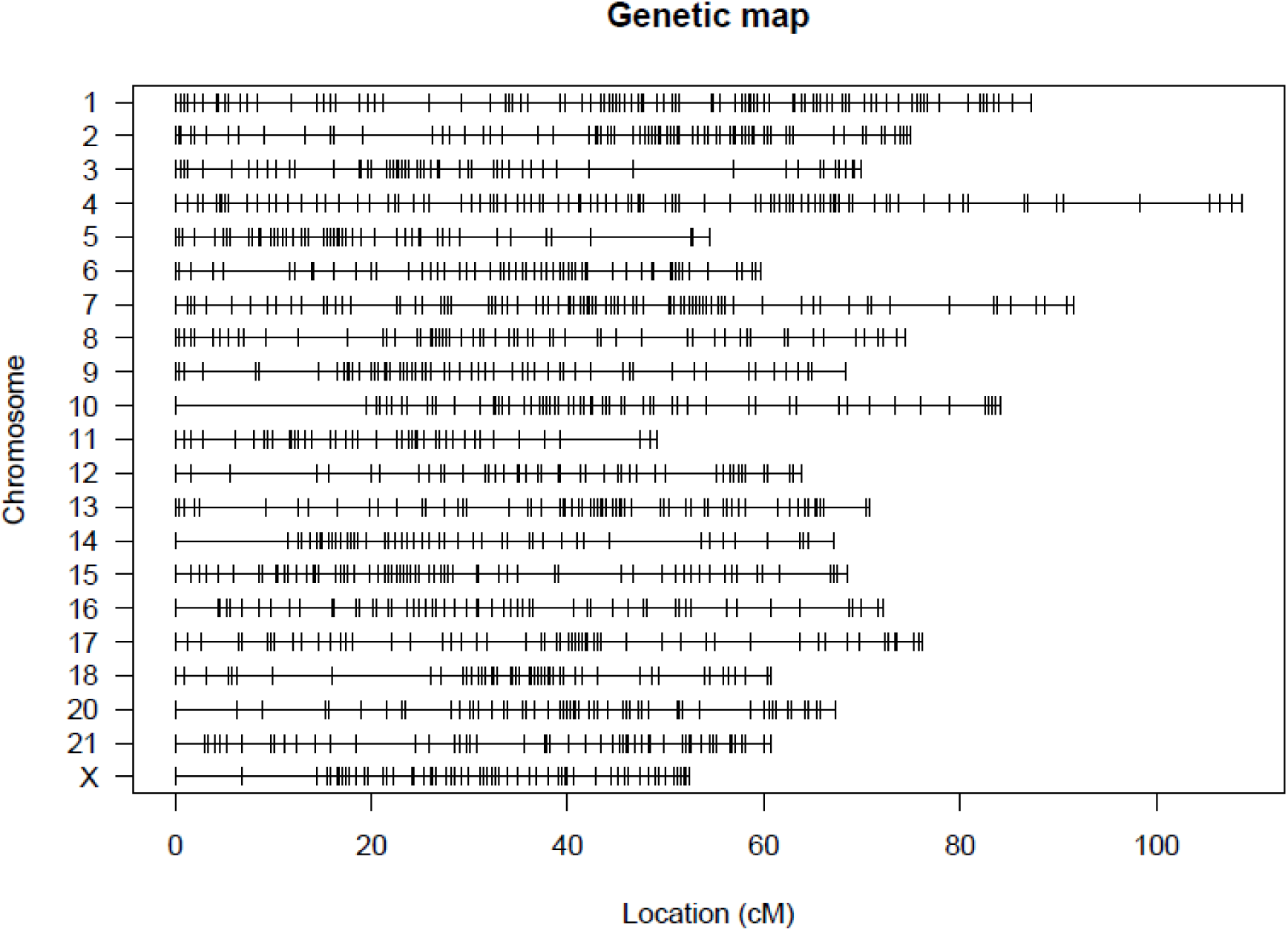
The stickleback linkage map consists of 21 linkage groups (20 autosomal, 1 sex) assembled from 1853 markers.

**Figure S2.**
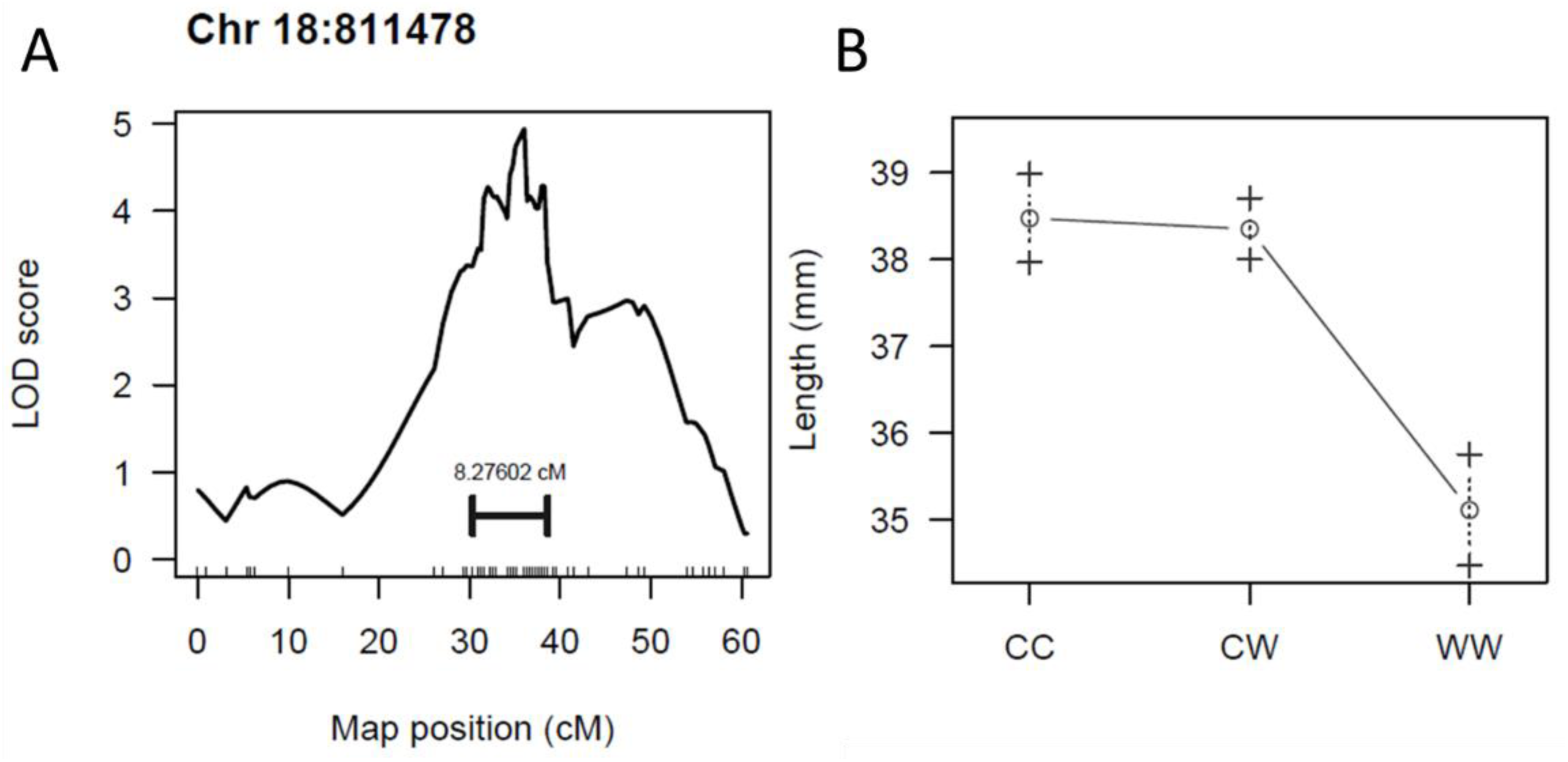
Body size (standard length) is mappable in F2 hybrids. (A) A QTL associated with body size was identified on chromosome 18, with the colored marker indicating the confidence interval. (B) The effect plot displays the phenotypic average for homozygous common (CC), homozygous white (WW) and heterozygous alleles at this locus in the F2s. Commons are larger than whites, and the common allele was associated with a larger size at this site.

**Figure S3.**
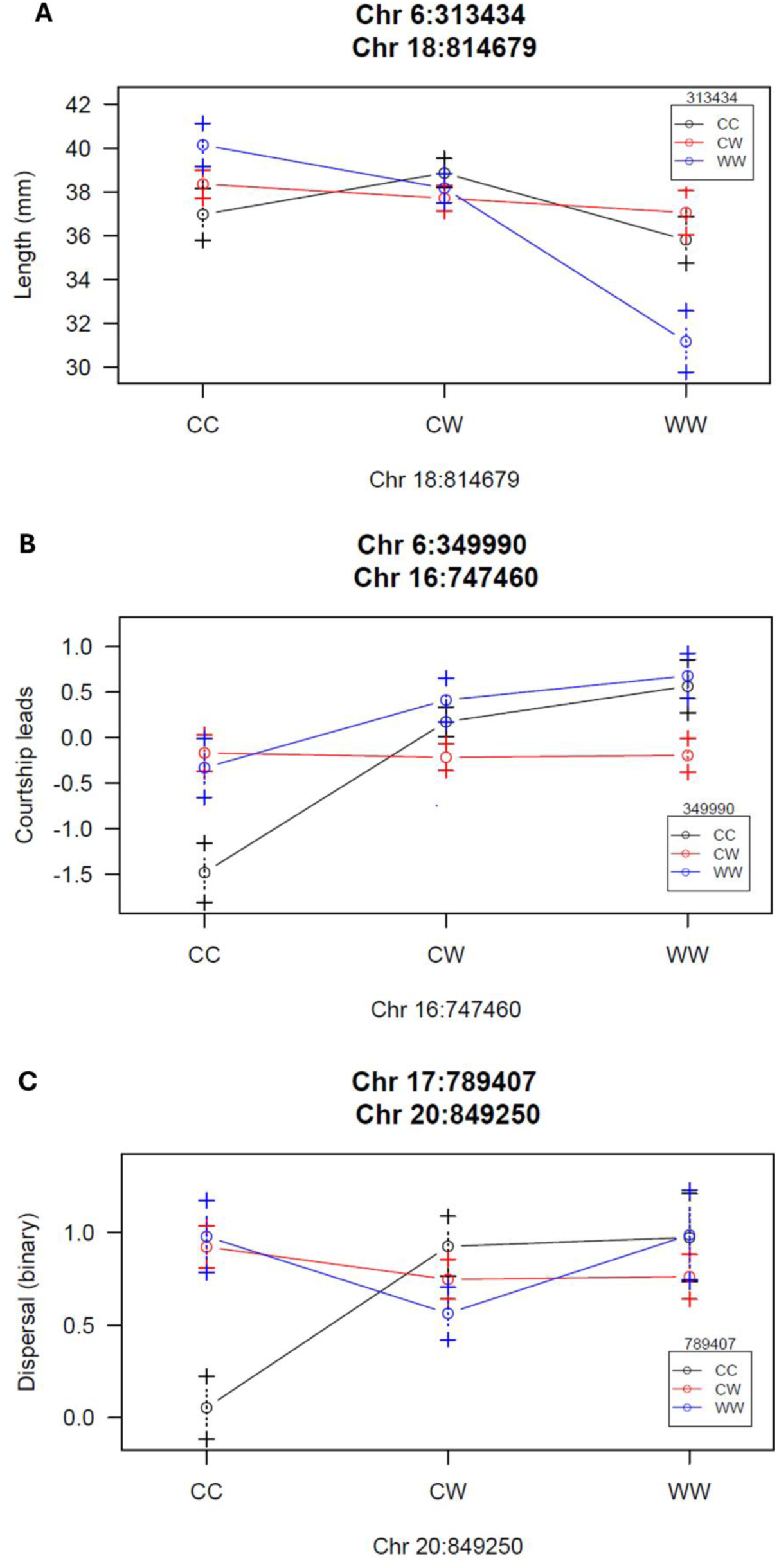
Interacting locus pairs from two-dimensional genome scans. Effect plots display the phenotypic averages for homozygous common (CC), homozygous white (WW) and heterozygous alleles at two sites. (A) Individuals with homozygous white alleles at both sites are significantly smaller in body size, matching patterns observed in natural common and white populations. (B) Individuals with homozygous common alleles at both sites exhibit significantly fewer leads during courtship, matching patterns observed in natural common and white populations. (C) Individuals with homozygous common alleles at both sites exhibit fewer egg dispersals.

**Figure S4.**
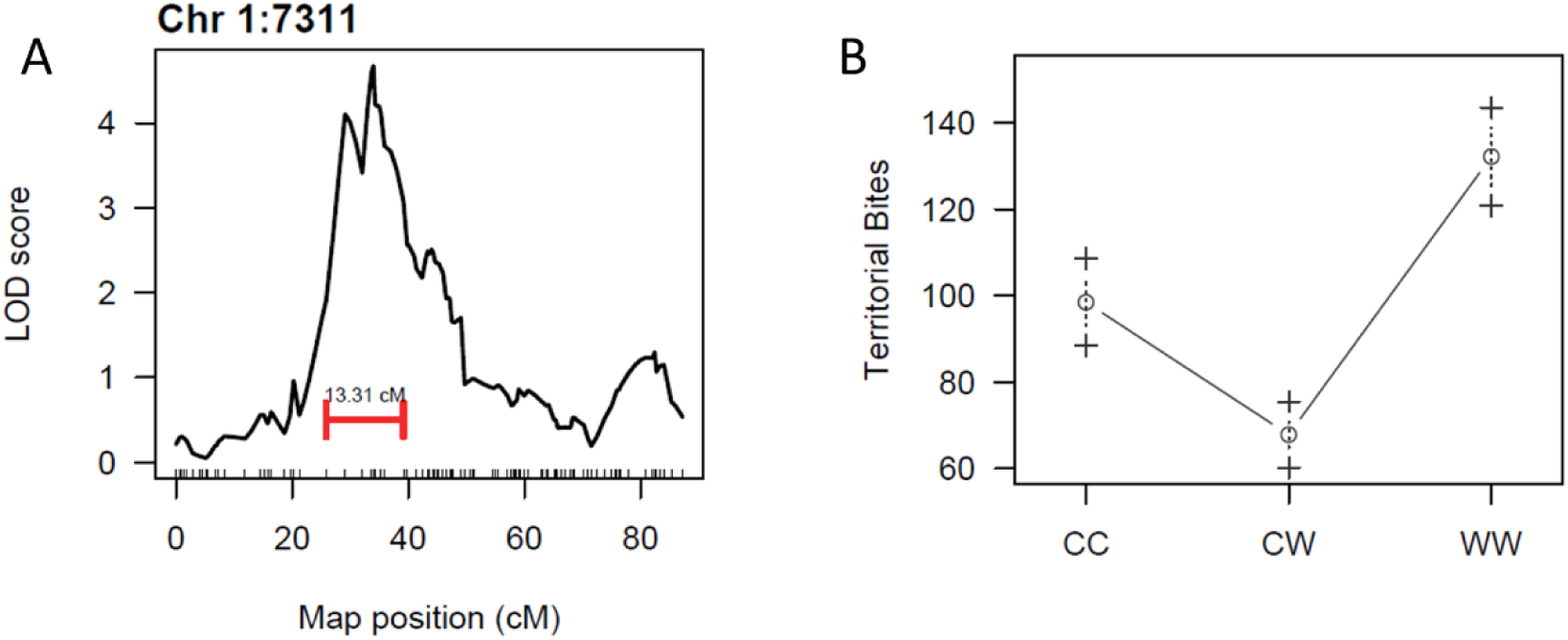
Territorial aggression is mappable in F2 hybrids. (A) A QTL associated with territorial bites was identified on chromosome 1, with the colored marker indicating the confidence interval. (B) The effect plot displays the phenotypic average for homozygous common (CC), homozygous white (WW) and heterozygous alleles at this locus in the F2s. Whites trend towards higher aggression than commons, and the white allele was associated with greater aggression at this site.

**Figure S5.**
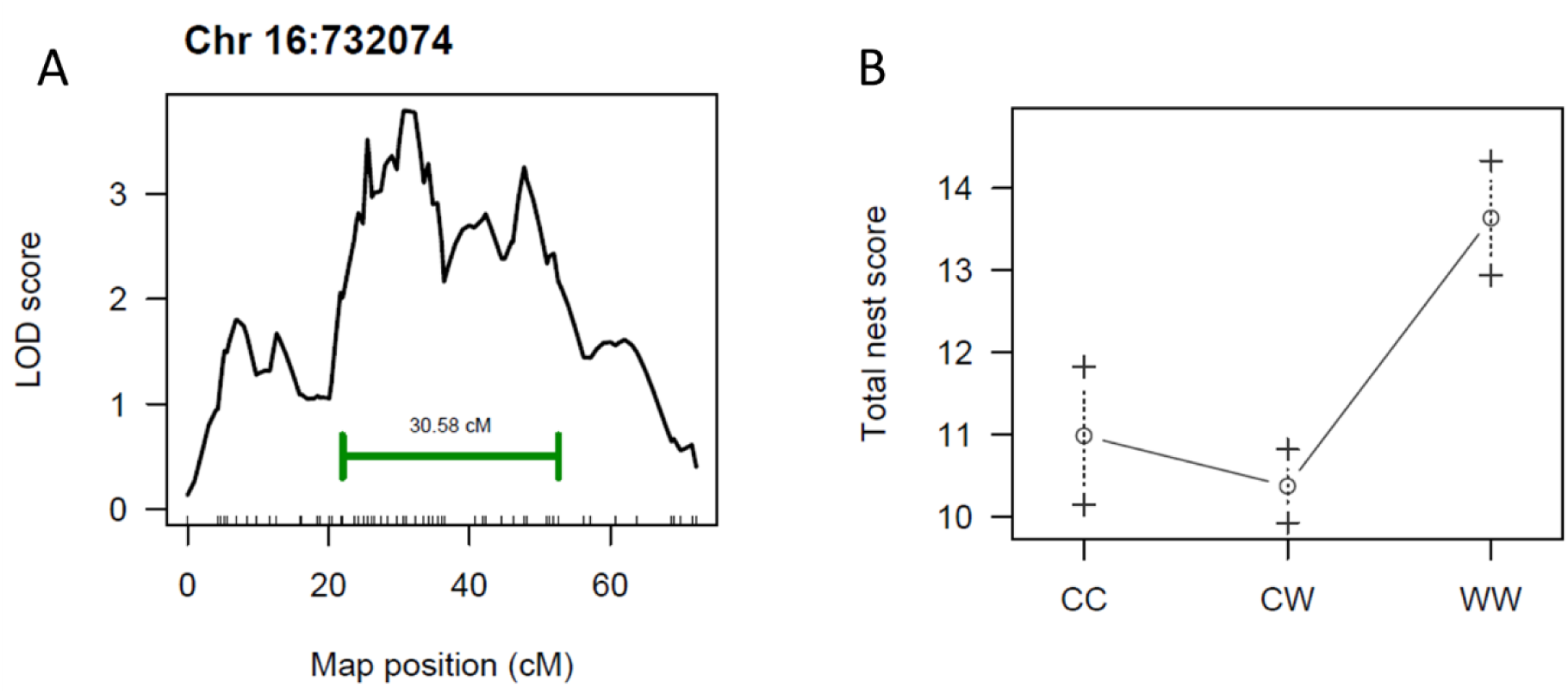
Nest architecture is mappable in F2 hybrids. (A) A QTL associated with nest score was identified on chromosome 16, with the colored marker indicating the confidence interval. (B) The effect plot displays the phenotypic average for homozygous common (CC), homozygous white (WW) and heterozygous alleles at this locus in the F2s. Whites build higher-scoring nests than commons, and the white allele was associated with higher-scoring nests at this site.

**Figure S6.**
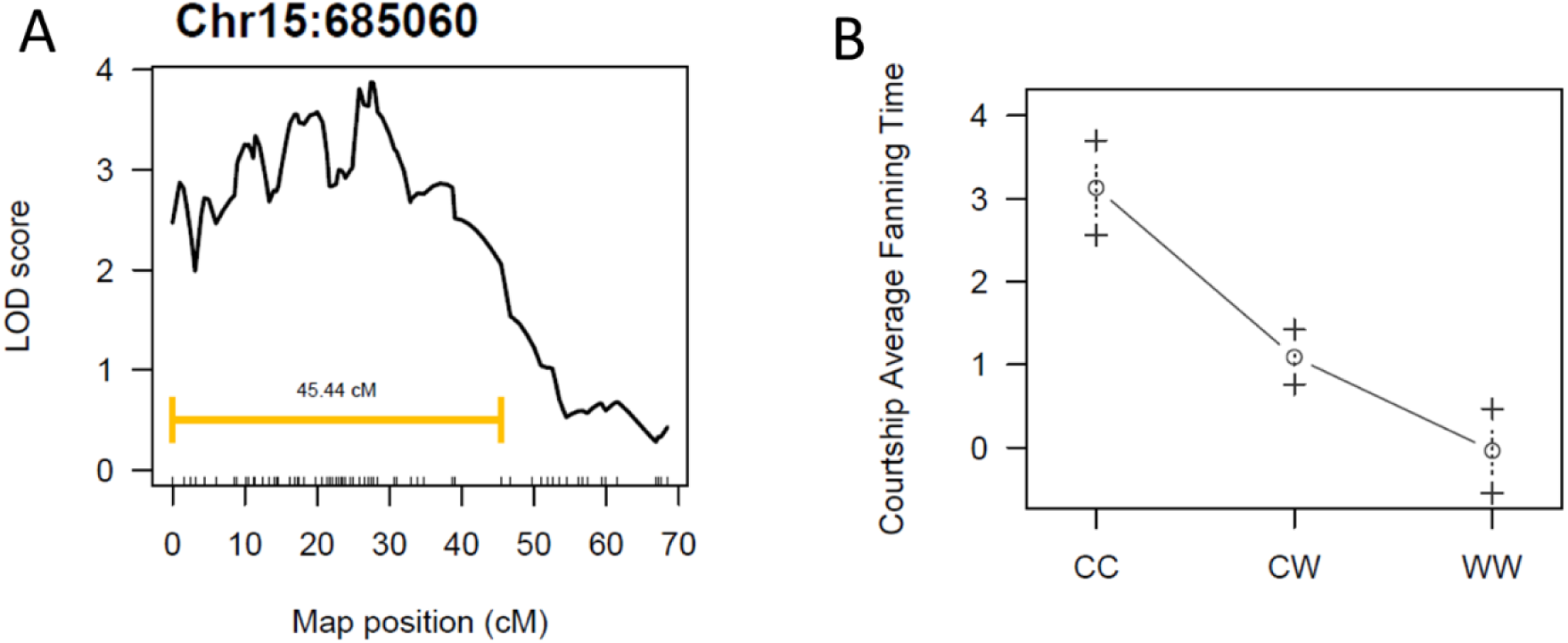
Courtship behaviors are mappable in F2 hybrids. (A) A QTL associated with fanning was identified on chromosome 15, with the colored marker indicating the confidence interval. (B) The effect plot displays the phenotypic average for homozygous common (CC), homozygous white (WW) and heterozygous alleles at this locus in the F2s. Commons exhibit greater fanning than whites, and the common allele was associated with more fanning at this site.

**Figure S7.**
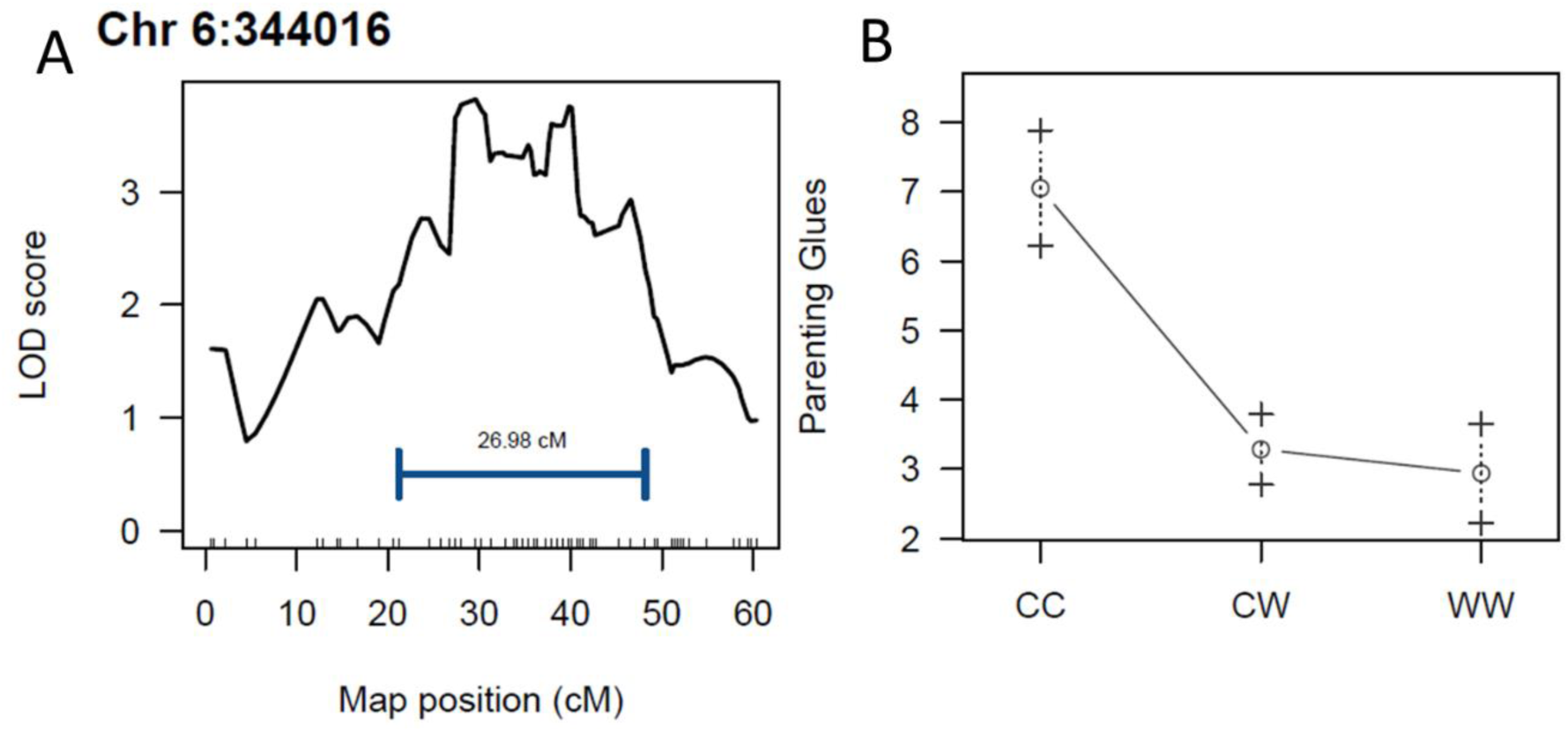
Parenting behaviors are mappable in F2 hybrids. (A) A QTL associated with glues during parenting was identified on chromosome 6, with the colored marker indicating the confidence interval. (B) The effect plot displays the phenotypic average for homozygous common (CC), homozygous white (WW) and heterozygous alleles at this locus in the F2s. Commons glue more frequently than whites, and the common allele was associated with greater gluing rates at this site.

**Figure S8.**
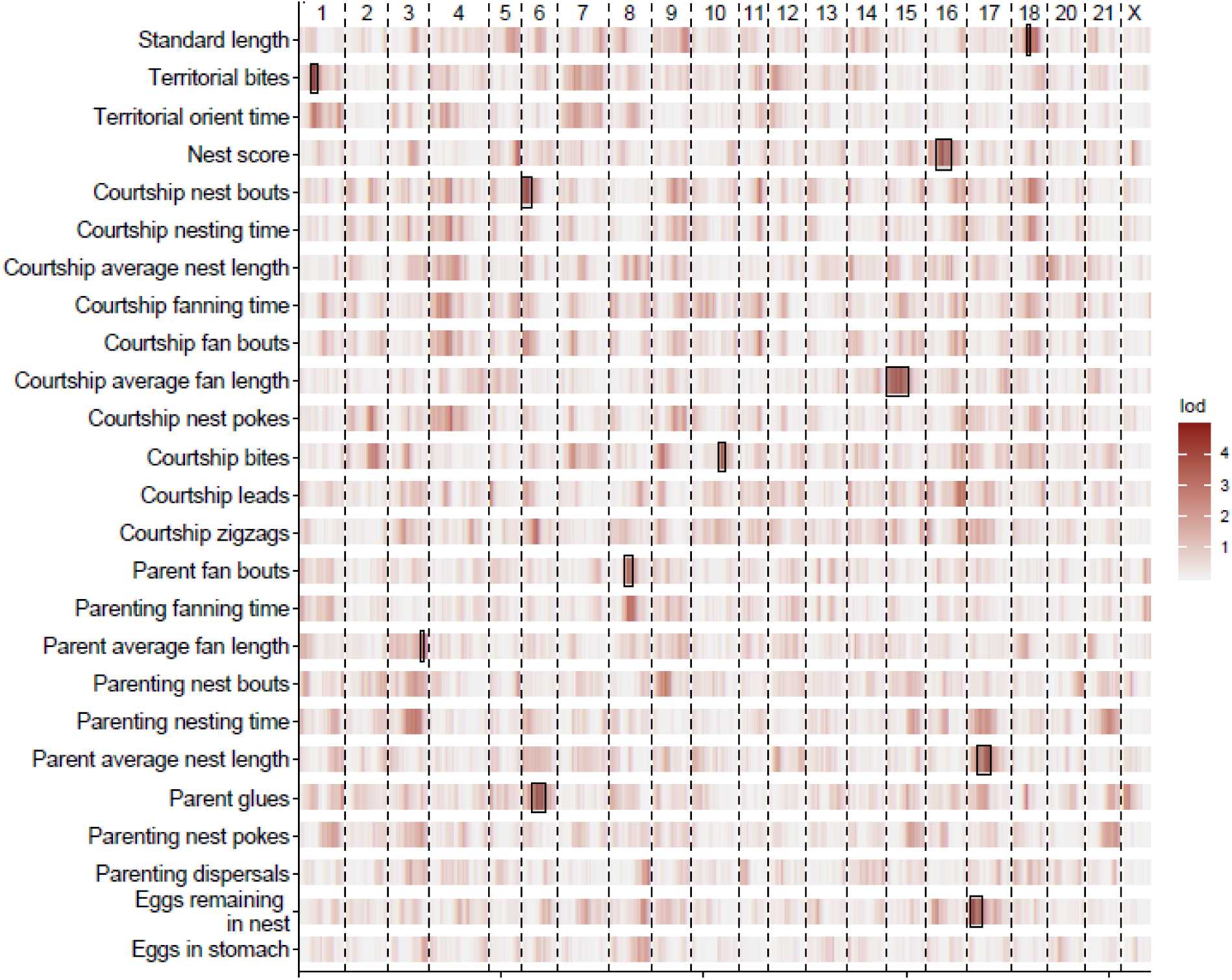
Mapping results of all analyzed reproductive traits. The LOD score for each trait is plotted across all chromosomes, with darker sections indicating higher LOD scores. Significant peaks and their confidence intervals are indicated with boxes. Chromosome 19 is the sex chromosome and is renamed here as “X”.

**Table S1.**
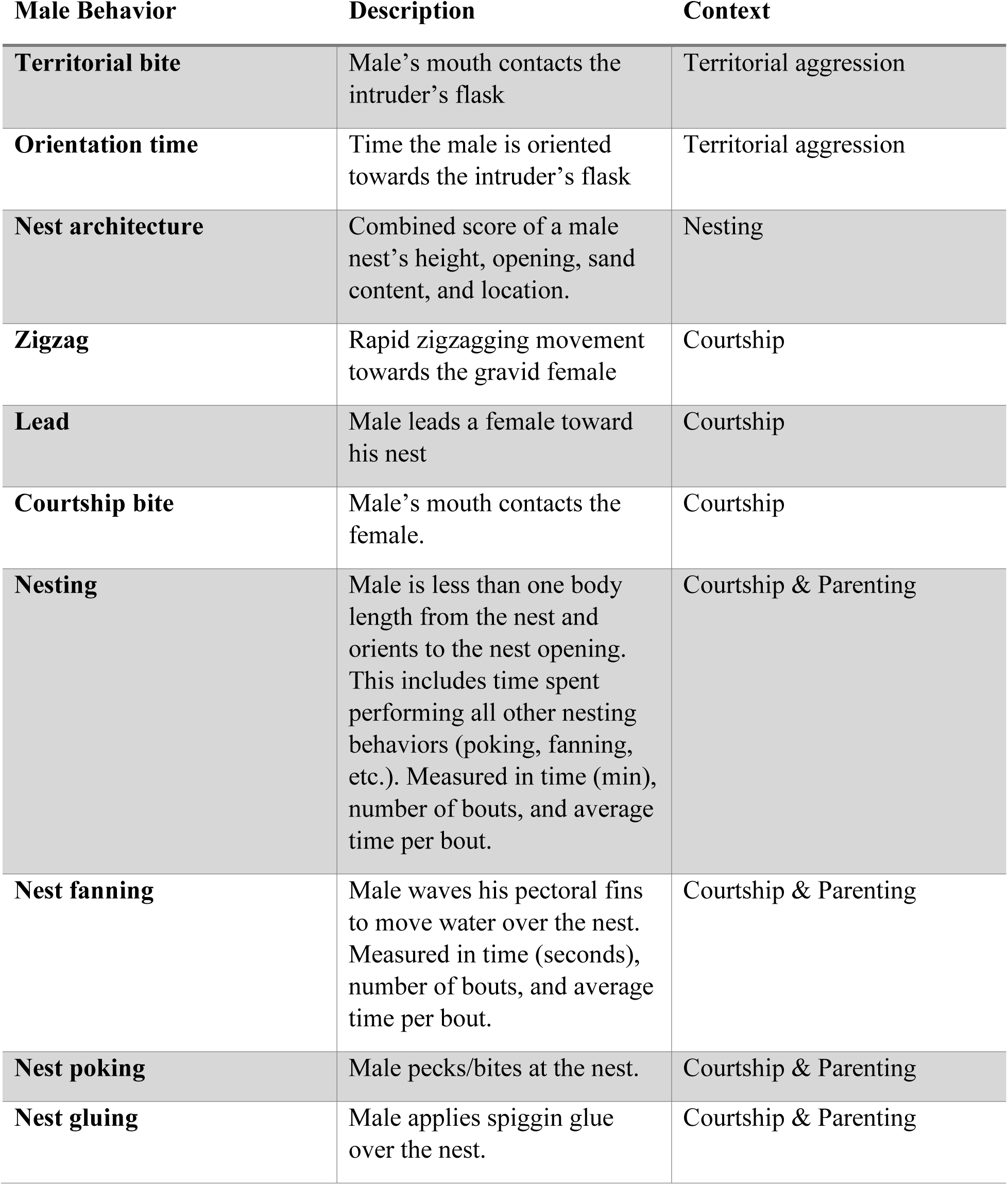
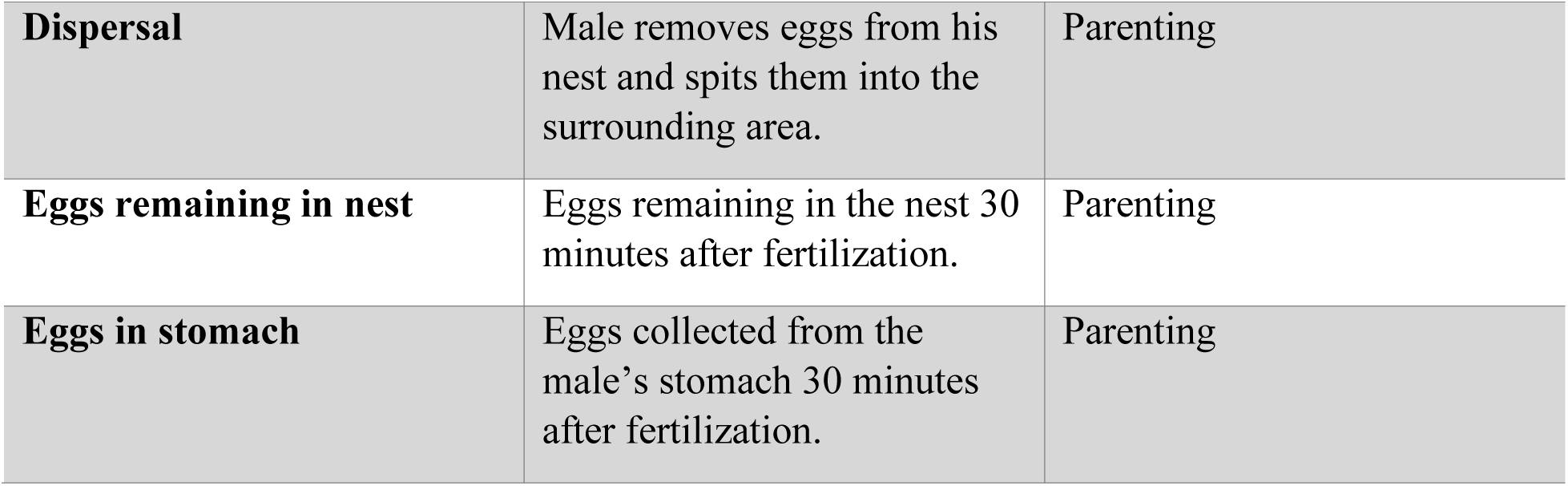
Male behaviors and their descriptions from territorial, nesting, courtship, and parenting measurements. Traits measured across courtship and parenting observations are analyzed separately.

| Male Behavior | Description | Context |
| --- | --- | --- |
| **Territorial bite** | Male’s mouth contacts the intruder’s flask | Territorial aggression |
| **Orientation time** | Time the male is oriented towards the intruder’s flask | Territorial aggression |
| **Nest architecture** | Combined score of a male nest’s height, opening, sand content, and location. | Nesting |
| **Zigzag** | Rapid zigzagging movement towards the gravid female | Courtship |
| **Lead** | Male leads a female toward his nest | Courtship |
| **Courtship bite** | Male’s mouth contacts the female. | Courtship |
| **Nesting** | Male is less than one body length from the nest and orients to the nest opening. This includes time spent performing all other nesting behaviors (poking, fanning, etc.). Measured in time (min), number of bouts, and average time per bout. | Courtship & Parenting |
| **Nest fanning** | Male waves his pectoral fins to move water over the nest. Measured in time (seconds), number of bouts, and average time per bout. | Courtship & Parenting |
| **Nest poking** | Male pecks/bites at the nest. | Courtship & Parenting |
| **Nest gluing** | Male applies spiggin glue over the nest. | Courtship & Parenting |
| **Dispersal** | Male removes eggs from his nest and spits them into the surrounding area. | Parenting |
| **Eggs remaining in nest** | Eggs remaining in the nest 30 minutes after fertilization. | Parenting |
| **Eggs in stomach** | Eggs collected from the male's stomach 30 minutes after fertilization. | Parenting |

**Table S2.** Marker distribution in the linkage map.

| Linkage Group | Number of markers | Length (cM) | Average marker spacing | Maximum marker spacing |
| --- | --- | --- | --- | --- |
| 1 | 146 | 87.236 | 0.601628 | 4.639 |
| 2 | 117 | 74.868 | 0.645414 | 7.131 |
| 3 | 78 | 69.922 | 0.908078 | 10.139 |
| 4 | 122 | 108.665 | 0.898058 | 7.866 |
| 5 | 62 | 54.406 | 0.891902 | 10.139 |
| 6 | 102 | 59.742 | 0.591505 | 6.767 |
| 7 | 134 | 91.538 | 0.688256 | 6.048 |
| 8 | 71 | 74.44 | 1.063429 | 4.988 |
| 9 | 80 | 68.353 | 0.865228 | 6.048 |
| 10 | 87 | 84.154 | 0.978535 | 19.543 |
| 11 | 62 | 49.029 | 0.803754 | 8.238 |
| 12 | 77 | 63.885 | 0.840592 | 8.825 |
| 13 | 99 | 70.716 | 0.721592 | 6.767 |
| 14 | 65 | 67.206 | 1.050094 | 11.526 |
| 15 | 88 | 68.437 | 0.786632 | 6.406 |
| 16 | 75 | 72.186 | 0.975486 | 4.957 |
| 17 | 92 | 76.174 | 0.837077 | 4.988 |
| 18 | 74 | 60.636 | 0.83063 | 10.139 |
| 20 | 95 | 67.331 | 0.716287 | 6.406 |
| 21 | 68 | 60.631 | 0.90494 | 6.048 |
| X | 59 | 52.442 | 0.904172 | 7.628 |
| overall | 1853 | 1481.997 | 0.80895 | 19.543 |

**Table S3.** Results from mixed models testing for an effect of female interest on male courtship behavior. Most traits were significantly associated with female interest level and were z-score normalized.

| Behavior | DenDF | F | P |
| --- | --- | --- | --- |
| Zigzags | 160.99 | 0.0709 | 0.932 |
| Bites | 118.29 | 0.9531 | 0.389 |
| Leads | 178.18 | 11.456 | **<0.001** |
| Pokes | 159.46 | 16.587 | **<0.001** |
| Glues | 174.98 | 23.94 | **<0.001** |
| Nesting time (min) | 190.95 | 49.296 | **<0.001** |
| Nesting bouts | 151.76 | 27.589 | **<0.001** |
| Fanning time (s) | 169.98 | 8.8074 | **<0.001** |
| Fanning bouts | 152.93 | 11.487 | **<0.001** |

